# COUPLING OF ENVIRONMENTAL AND DIRECT TRANSMISSION MECHANISMS: ANALYSIS OF A SIMPLE MODEL

**DOI:** 10.64898/2026.01.20.700625

**Authors:** José Manuel Islas, Baltazar Espinoza, Jorge X. Velasco-Hernández

## Abstract

We study an extension of an environmentally mediated epidemiological model that incorporates direct human-to-human transmission. While the original formulation accounted for environmental exposure, it did not include direct transmission between individuals. Allowing both transmission routes to interact leads to significant qualitative changes in the system dynamics. The analysis reveals multiple dynamical regimes governed by environmental and combined threshold quantities. The stability of the disease-free equilibrium is controlled by an environmental threshold, whereas a combined reproduction number determines the onset of multistability. For certain parameter ranges, endemic equilibria coexist with the disease-free equilibrium, giving rise to backward-type bifurcation behavior and sensitivity to initial conditions. Moreover, the direct transmission rate acts as an organizing parameter by inducing the emergence of an environmental-free equilibrium when exceeding its classical threshold. These results highlight how environmentally coupled transmission mechanisms can generate rich dynamics in low-dimensional models.

## 1 Introduction

Modern societies constantly face infectious disease threats and risk of contagion that challenge public health systems. These threats arise not only from direct human-to-human transmission but also through direct exposure to animal vectors and interaction with environmental sources such as contaminated water, food, air, and surfaces. The diversity of these transmission pathways highlights the need for integrated modeling approaches that address the complexity of disease prevention, surveillance, and control. Such approaches should explicitly account for human behavior, ecological change, and environmental vulnerability [6, 4, 2, 8]. In particular, integrated frameworks allow to examine the complete transmission loop, in which environmental reservoirs can sustain infection and generate complex threshold dynamics arising from source–sink processes often overlooked in models without explicit environmental coupling.

Risk management through disease control and prevention is essential for public health policies. However, the actual effects of intervention measures must be adequately evaluated to prevent unintended consequences. It is well known that implementing public health measures to reduce the impact of infectious agents in a population can potentially lead to subthreshold dynamics [15, 26, 3]. Mass vaccination campaigns, behavioral changes, and non-pharmaceutical interventions can alter epidemic dynamics and the sharp threshold for outbreak emergence, which is typically defined by the basic reproduction number [5, 9, 8, 7]. In such scenarios, an epidemic may persist even when the basic reproduction number ( ℛ_0_) falls below the critical threshold for outbreak emergence. Subthreshold dynamics in epidemic models has drawn the attention of researchers during the last decade due to its important implications in public health and the mechanisms leading to subthreshold dynamics have been widely studied, as understanding these processes is crucial for developing strategies to control outbreaks [23, 25, 24, 12, 10, 22]. If a backward bifurcation exists, a transcritical bifurcation where the branch of endemic equilibrium arises with a negative slope, makes control impossible since it now depends on the initial condition; if, particularly the magnitude of the infected component that enters the population is large the system could stabilize itself at the endemic equilibrium below the reproductive number threshold. There are many mechanisms associated with the existence of backward bifurcations in directly transmitted diseases [10], while subthreshold dynamics have been documented in epidemic models studying communicable diseases, vector-borne transmitted diseases, and water-borne diseases [24].

The focus of this work is to address a critical gap in traditional epidemiological models, which often consider only a single mode of transmission or treat environmental effects as exogenous processes. By explicitly integrating environmental reservoirs into the transmission process, we aim to understand how different modes of transmission interact with a population and shape key epidemiological outcomes, such as prevalence levels and potential for disease persistence. This explicit coupling allows us to disentangle complex disease persistence conditions and the relative contributions of direct and environmental transmission pathways for scenarios where outbreaks evolve in settings where sanitation, climate conditions, or pathogen durability play significant roles. In this study, we use a mathematical model that incorporates two transmission pathways. The first is direct transmission between hosts, which essentially follows a standard Susceptible-Infected structure. The second is an environmental transmission pathway associated with a polluted space that contains pathogens and evolves dynamically according to a logistic-type growth law, driven, for example, by bacterial communities [27, 14]. The inclusion of logistic growth implies that the carrying capacity for the pathogen is bounded, which limits the number of pathogens that would be present in this environment. Environmentally transmitted diseases are challenging to model because of the difficulty to measure the abundance of a pathogen in a solution, either in air, water, or any other environmental component that can produce disease when in contact with humans [13, 16]. In our model, we assume that the susceptible population is infected by direct contact with an infectious individual and through contact with either fomites or other vectors in a polluted environment, such as drinking water, for example. The coupling with the environment comes through the shedding of pathogens by the infectious individuals that find their way into the environment; moreover, we assume that the polluted environment has an external input independent of the number of infected individuals in the populations where we are modeling our process. For previous approach with different emphasis on this problem, see [18].

## 2 Model formulation

We propose coupling the susceptible–infected model for human-to-human transmission with the metapopulation model introduced in [17], thereby accounting for environmental transmission:

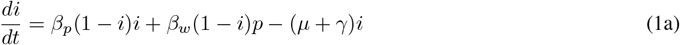

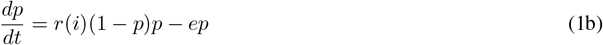

where *β*_*p*_ represents the human transmission rate, *β*_*w*_ represents the environment transmission rate, *r*(*i*) is the environment recruitment rate as a function of the infected population, and *b* = *µ* producing a constant population size, that is *s* + *i* = 1. For simplicity, we assume a baseline recruitment rate *r*_0_ for environmental infection. Moreover, we assume that in the presence of infectious individuals, the environmental infection rate linearly increases as *ai*, where *a* is a constant rate of pathogen shedding into the environment. As a result, the environment inflow rate is assumed to have the form *r*(*i*) = *r*_0_ + *ai*. This model is a minimal version of the model in [14] where similar behavior to the one explored here appears, albeit in a more complex context.

The dynamics of the decoupled models (1a) or (1b) is well known (see [11, 17]): in the absence of infection, the dynamics of model (1b) is determined by the environmental threshold parameter

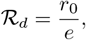

which depends only on the environmental pollution rates, and we can note that

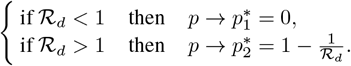

On the other hand, the dynamics of the host-only model (1a) is determined by the basic reproduction number

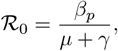

represents the average number of new infections generated by a single infectious individual. We have that

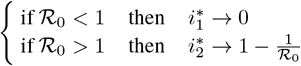

### 2.1 The coupled model undergoes a double forward bifurcation

When the systems are coupled, we begin by examining the conditions for a positive equilibrium in the environment. Using the linear recruitment function *r*_0_ + *ai*^∗^ and assuming that infected hosts have already reached an steady state, we obtain

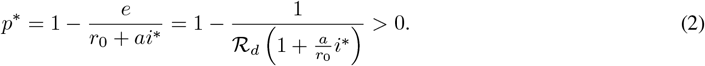

From (2), we conclude that the environment can sustain a positive equilibrium

i. **(Levins’ metapopulation dynamics)** when there are no host infections (*i*^∗^ = 0) and ℛ_*d*_ > 1
ii. **(Coupled dynamics)** ℛ_*d*_ < 1 but disease prevalence in the hosts is large enough (*i*^∗^ *> i*^+^) to compensate for the environment’s pathogen clearing rate. In this scenario, the minimum host prevalence necessary to sustain a positive environmental pollution level is

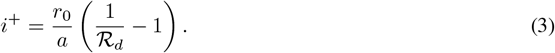

Alternatively, we can write, from (1a), the reproduction number ℛ that can sustain this environment endemic state as a function of *i*^+^:

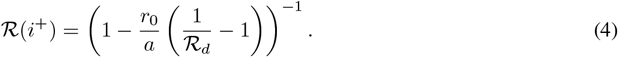

Therefore, there is a minimal prevalence level *i*^+^, such that the associated shedding of pathogens by these infected hosts can produce an environmental positive equilibrium state in an environment unable to support one by itself ( ℛ_*d*_ < 1). In fact, we can calculate the minimum ℛ_*d*_ associated with this *i*^+^ < 1, for which sustained host infections can induce the environment to reach an equlibrium state *p*^∗^. In other words if

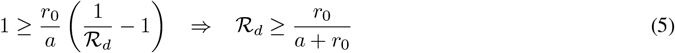

Consequently, if 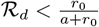 there is no host prevalence level *i*^+^ that can drive the environment to apositive equilibrium state.

In summary, for the coupled model, the environment can support an endemic state *p*^∗^ > 0 whenever

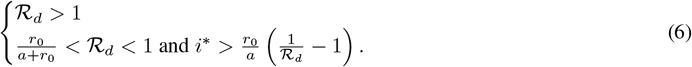

Figure 1 illustrates condition (6). Initially, the classical transcritical bifurcation driven by host-host transmission occurs at ℛ_0_ = 1. Moreover, once the host endemic equilibrium surpasses the critical threshold *i*^+^, the environmental transmission undergoes a transcritical bifurcation (*p*^∗^ > 0). In these scenarios, the endemic equilibrium on hosts shows a second transcritical bifurcation driven by a double source of infections (direct by hosts and indirect through environmental infection), ultimately increasing the system’s total prevalence. Note that the second transcritical bifurcation occurs when ℛ_0_ > 1, *i*^∗^ *> i*^+^, and 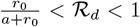. Figure 1(a) shows the classic forward bifurcation on the infectious individual’s dynamics in the absence of environmental pollution, for *i*^∗^ *< i*^+^; and the increased prevalence of infectious host population due to the existence of environmental contamination, for *i*^∗^ *> i*^+^. Figure 1(b) shows the dynamics of the environmental component when ℛ_*d*_ = 0.6, as a function of ℛ_0_. For the selected simulations, it is required that ℛ_0_ > 1.4 to lead the environment to an endemic state.

**Figure 1.**
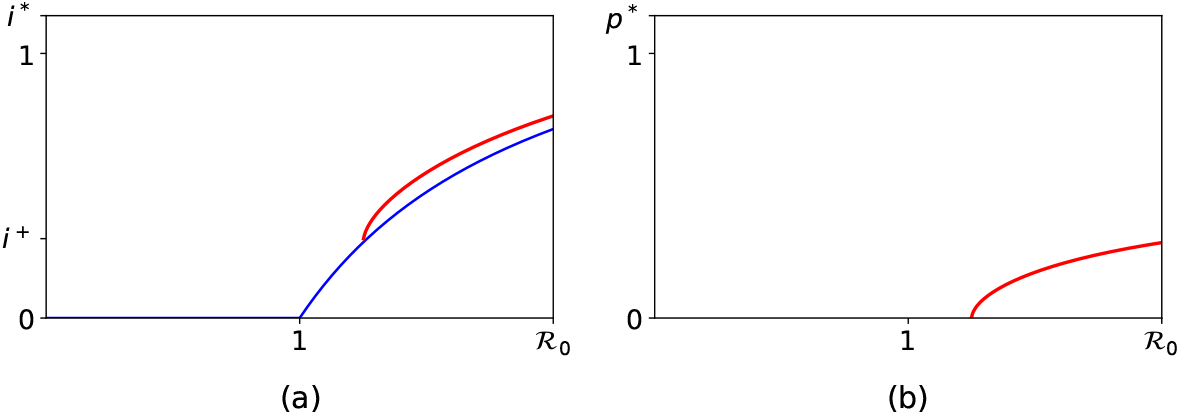
The double transcritical bifurcation. The host’ prevalence endemic state larger than the threshold *i*^+^ leads the dynamics of the environment to undergo a transcritical bifurcation, which produces a second transcritical bifurcation, this time on the host infection dynamics. Our simulations assume the following parameters ℛ_*d*_ 0.85, *β*_*w*_ = 0.4, *µ* = *γ* = 0.5, *e* = 0.47, *a* = 0.35, and *r*_0_ = 0.4, where *i*^+^ = 0.285.

## 3 Multistability dynamics in the coupled model

Figure 2 shows that the endemic state in the coupled system can be maintained even when the hosts’ (ℛ _0_) and environment’s (ℛ _*d*_) reproductive numbers are less than one. It is interesting to note that the environment’s backward bifurcation dominates the dynamics of the coupled system.

**Figure 2.**
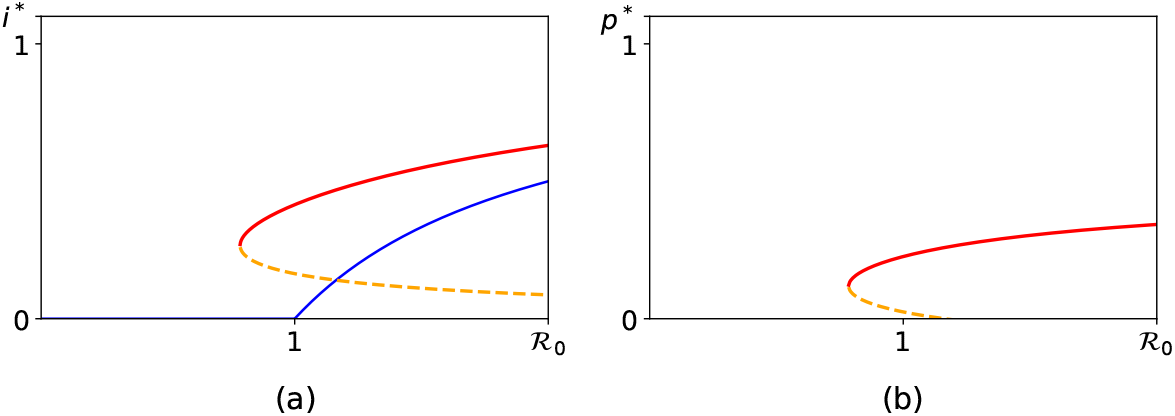
Infectious individuals’ and environment’s endemic levels are supported when ℛ_0_ < 1 and ℛ_*d*_ ≈ 0.85. Parameters set: *β*_*w*_ = 1.3, *β*_*p*_ ∈ [0.6, 1], *µ* = *γ* = 0.5, *e* = 0.47, *a* = 0.5, *r*_0_ = 0.4.

### 3.1 General results on the coupled system

In this section, we present a formal analysis of the coupled model (1a)-(1b). The proofs of the propositions, along with illustrative figures, are provided in Appendix A. We begin by verifying that the model is well-defined within the domain of biological interest.

#### Proposition 1

*Let* Ω = {(*i, p*) | 0 ≤ *i* ≤ 1, 0 ≤ *p* ≤ 1}. *Then* Ω *is an invariant set for model (1a)-(1b)*.

To analyze the model’s dynamics, we first identify its fixed points. Accordingly, we set *r*_0_ = *R*_*d*_ *e*, and solve the following system of algebraic equations:

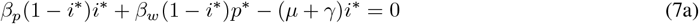

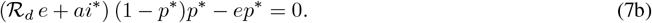

First, from equation (7a), we obtain

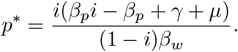

Note that for the feasibility of the equilibria (*i*^∗^, *p*^∗^) ∈ Ω, we have

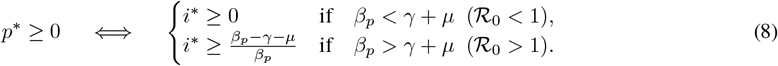

Next, substituting *p*^∗^ in equation (7b) we obtain the polynomial of degree 5

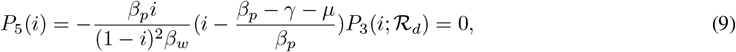

Where

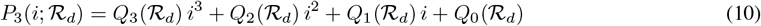

With

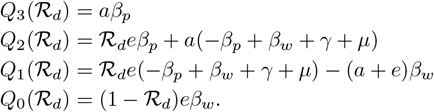

#### 3.1.1 DFE and EFE Equilibria

From (9), two trivial solutions of (7a)–(7b) can be identified: the disease-free equilibrium DFE = (0, 0) and the environmental pollution-free equilibrium EFE = (*i*_EFE_, 0), where

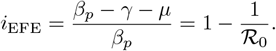

The latter is biologically meaningful only when *β*_*p*_ *> γ* + *µ* (ℛ_0_ > 1). Furthermore, it is immediate to verify that when *β*_*p*_ = *γ* + *µ* (ℛ_0_ = 1), these two equilibria coincide. We may therefore state the following proposition:

##### Proposition 2

*The model (1a)-(1b) experiences a transcritical bifurcation at* DFE *when β*_*p*_ = *γ* + *µ*.

Note that Proposition 2 establishes the appearance of EFE as an equilibrium in Ω once *β*_*p*_ crosses the bifurcation threshold *γ* + *µ*. Moreover, this equilibrium arises with local asymptotic stability, as shown in the following result.

##### Proposition 3

*Consider the model (1a)-(1b):*

a. *If β*_*p*_ *< γ* + *µ and* ℛ_*d*_ < 1, DFE *is locally asymptotically stable and* EFE *is unstable*.
b. *If β*_*p*_ *> γ* + *µ and* 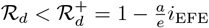 DFE *is unstable and* EFE *is locally asymptotically stable*.
c. *Otherwise, both equilibria are unstable*.

The above analysis shows that when only the DFE is feasible, it is stable for ℛ_*d*_ < 1 and unstable for ℛ_*d*_ > 1. In contrast, once the EFE becomes feasible, it is stable for 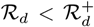 and unstable for 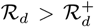, while the DFE remains unstable for all values of ℛ_*d*_. We remark that as ℛ_*d*_ depends exclusively on environmental parameters, it does not represent the average number of secondary transmissions from one infected person, as in the standard way of reproduction number i.e. it is not a reproductive number [21].

#### 3.1.2 Endemic Equilibria

To determine the conditions under which multiple feasible endemic equilibria arise, we examine the third-degree polynomial (10). Observe that *Q*_3_(ℛ_*d*_) > 0 for all ℛ_*d*_, and that the inequality *Q*_2_(ℛ_*d*_) < 0 necessarily implies *Q*_1_(ℛ_*d*_) < 0. Applying Descartes’ rule of signs, we conclude that the case *Q*_1_(ℛ_*d*_) < 0 is the only scenario that permits either two or zero positive roots when *Q*_0_(ℛ_*d*_) > 0 (corresponding to ℛ_*d*_ < 1), and yields a single positive root when *Q*_0_(ℛ_*d*_) < 0 (i.e., ℛ_*d*_ > 1).

As observed, ℛ_*d*_ = 1 constitutes a critical threshold where the number of positive roots changes. In particular, the condition *Q*_1_(1) < 0 identifies an interval to the left of ℛ_*d*_ = 1 in which (10) admits two positive roots. Specifically, observe that the inequality *Q*_1_(1) < 0 reduces to

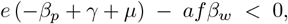

or, equivalently,

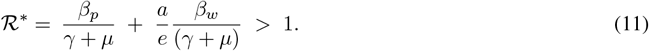

Note that the term 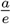 represents the average environmental pathogen load contributed by a single infected individual. The quantity 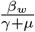denotes the average number of new infections generated by one unit of environmental pathogen over the duration of the infectious period. Their product,

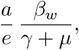

thus gives the expected number of new infections produced indirectly through the environment by a single infected individual. Therefore, the parameter ℛ^∗^ represents the expected number of secondary infections produced by an infected individual over its entire infectious period, accounting simultaneously for both transmission routes: direct and environmental.

**Remark 1** *Note from (11) that i*_EFE_ > 0 *implies* ℛ^∗^ > 1; *in other words, the feasibility of the* EFE *guarantees the possibility that (10) admits two positive roots. However, it is also possible to have* ℛ^∗^ > 1 *while i*_EFE_ < 0, *meaning that (10) may possess two positive roots even in the absence of a feasible* EFE. *Conversely, if* ℛ^∗^ < 1, *the* EFE *is unfeasible*.

Once ℛ^∗^ > 1, determining the precise interval in which (10) admits two positive roots requires computing its discriminant, given by

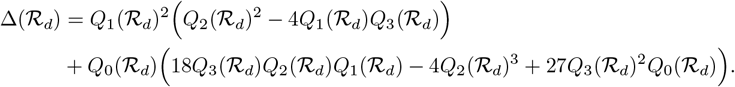

If Δ(ℛ _*d*_) > 0, then (10) has three real roots, whereas if Δ(ℛ _*d*_) < 0, it has only one real root. Note that, since *Q*_0_(1) = 0, we necessarily have Δ(1) > 0. Therefore, because the only scenario in which (10) admits two positive roots occurs for ℛ_*d*_ < 1, the interval supporting two positive roots must be of the form 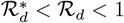, where 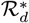 is such that 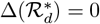. We summarize the results above as follows.

##### Proposition 4

*Consider the polynomial (10), with R*_*d*_ *and* 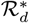 *defined above:*

a. *If* ℛ_*d*_ > 1 *then (10) has a unique positive root*.
b. *If* ℛ^∗^ > 1 *and* 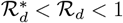 *then (10) has exactly two positive roots*.
c. *Otherwise, (10) does not have positive roots*.

Proposition 4 identifies the parameter regimes in which the polynomial (10) possesses two positive roots. However, these roots do not necessarily correspond to feasible endemic equilibria of the full model (1a)–(1b). According to the feasibility condition (8), when *β*_*p*_ *< γ* + *µ*, an equilibrium (*i*^∗^, *p*^∗^) is biologically admissible provided that *i*^∗^ > 0. In contrast, when *β*_*p*_ *> γ* + *µ*, the condition *i*^∗^ > 0 is no longer sufficient: feasibility additionally requires *i*^∗^≥*i*_EFE_.In other words, the appearance of the EFE equilibrium can restrict or prevent the existence of endemic equilibria. To analyze this interaction, we now study the transcritical bifurcations that arise in the model, as summarized in the following result.

##### Proposition 5

a. *The model (1a)-(1b) experiences a transcritical bifurcation at the* DFE *as the parameter varies through the bifurcation value* ℛ_*d*_ = 1.
b. *The model (1a)-(1b) experiences a transcritical bifurcation at the* EFE *as the parameter varies through the bifurcation value* 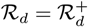.

Proposition 5 establishes the existence of two transcritical bifurcations in the model (1a)–(1b), which, in addition, may be accompanied by a change in the stability of the DFE and EFE. Figure 3 illustrates these bifurcations. First, we note that if *β*_*p*_ *< γ* + *µ*, the bifurcation at the EFE will not occur within Ω, and thus only the transcritical bifurcation at the DFE is observed, which in this case is the only one that modifies the stability of the system. Figures 3a and 3b show the bifurcation in this scenario when ℛ*d <* 1 and ℛ*d >* 1, respectively. On the other hand, if *β*_*p*_ *> γ* + *µ*, the transcritical bifurcation that occurs at the EFE determines the existence of endemic equilibria, since whenever the EFE is feasible, any equilibrium with *i*^∗^ *< i*_*EF E*_ is not feasible in Ω. Figures 3c and 3d show the transcritical bifurcation at the EFE in this scenario; the green continuous curve represents the endemic equilibrium of the system, whereas the green dotted curve represents the positive roots *i*^∗^ of (10), which satisfy *i*^∗^ *< i*_EFE_ and are therefore not feasible. Besides, we can observe, as mentioned in Proposition 5, that the DFE is unstable for all ℛ_*d*_. Moreover, note that as *i*_EFE_ increases, points on the bifurcation curve with *i*^∗^ *< i*_EFE_ become unfeasible; however, in Figure 3c, we see that there are still two positive roots to the left of ℛ_*d*_ < 1, but not in the interval 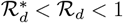. Such points will exist for 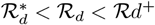. Finally, Figure 3d shows that, for sufficiently large values of *i*_EFE_, it will no longer be possible to observe two endemic equilibria for ℛ_*d*_ < 1; in this case, only a single positive root is present for 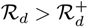.

**Figure 3.**
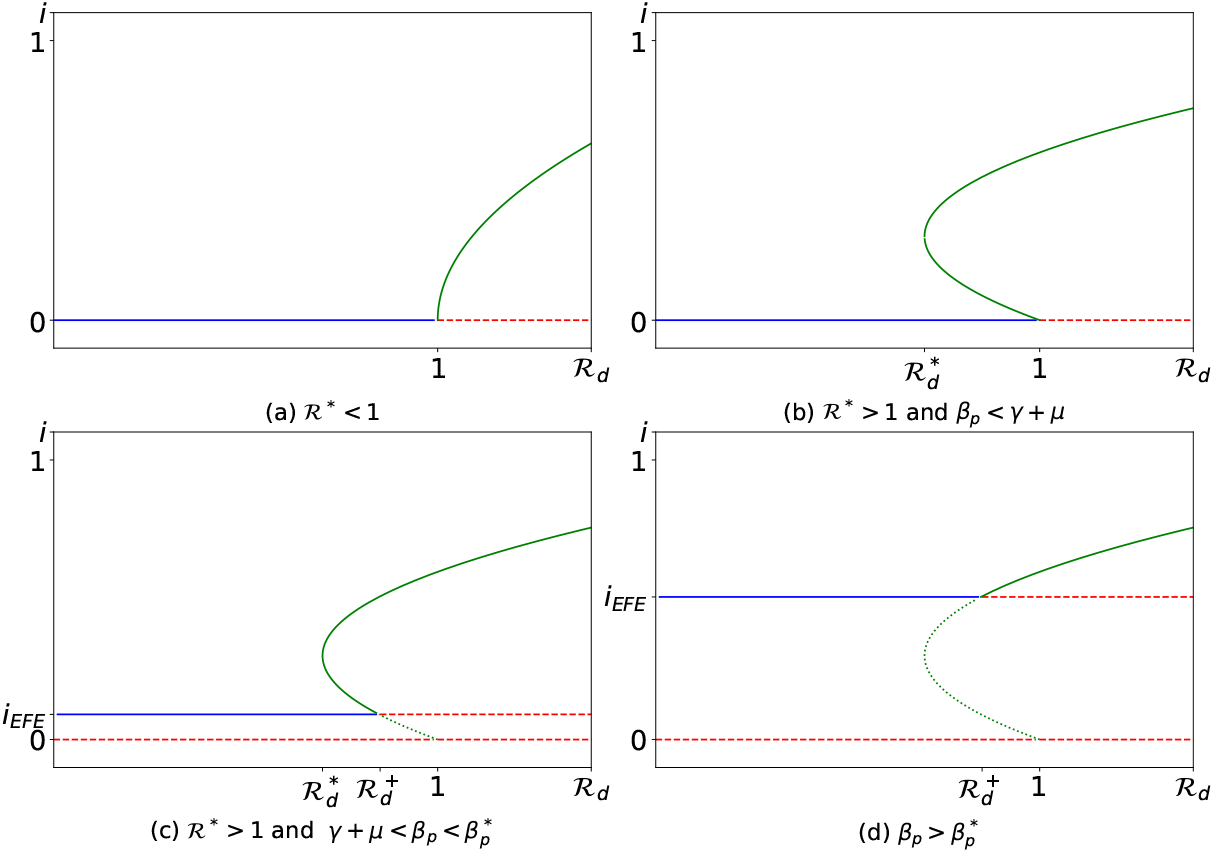
Interaction of transcritical bifurcations of Proposition 5 with the roots of polynomial (10) from Proposition 4. Green continuous curves represent feasible positive roots, green dotted curves represent unfeasible positive roots, blue represent stable equilibrium and red unstable equilibrium

The above suggests that there is a maximum value 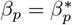 for which two endemic equilibria can exist in the model (1a)–(1b) even when ℛ_*_> 1. When 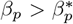, one or both of the potential equilibria may satisfy *i*^∗^ *< i*_EFE_, which makes them unfeasible in Ω. This threshold is given by

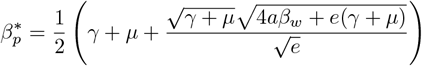

and allows us to classify the number of endemic equilibria in the following way.

##### Proposition 6

*Consider the model (1a)-(1b)*

a. *If* ℛ_∗_ < 1, *the model has a unique endemic equilibrium for ℛ*_*d*_ > 1.
b. *If* ℛ_∗_ > 1 *and*

1. *β*_*p*_ *< γ* + *µ, there exist exactly two endemic equilibria for*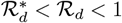, *and one endemic equilibrium for ℛ*_*d*_ > 1,
2. 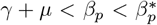, *there exist exactly two endemic equilibria for*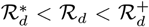, *and one endemicequilibrium for* 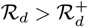,
3. 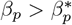, *there exists a unique endemic equilibrium for* 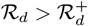.

Next, in order to characterize the stability of the endemic equilibria of model (1a)-(1b), we establish the following propositions, which identifies the conditions under which such equilibria persist or lose stability and thus clarifies the dynamical structure of the system.

##### Proposition 7

*If* (*i*^∗^, *p*^∗^) *is an endemic equilibrium, that is, if i*^∗^ *> i*_EFE_, *then the sign of the slope of the bifurcation curve, i vs* ℛ_*d*_ *of the model (1a)-(1b) in such point is determined by de sign of*

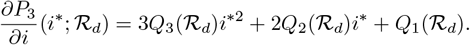

*If* 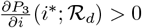, *the slope of the curve is positive and, if* 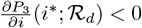, *the slope of the curve is negative*.

Therefore, we establish the stability of the endemic equilibrium based on its position on the bifurcation curve, as follows.

##### Proposition 8

*If an endemic equilibrium is located where the bifurcation curve has positive slope, then it is locally asymptotically stable, whereas if it is where the bifurcation curve has negative slope, it is unstable*.

### 3.2 Backward bifurcation

Now, in order to present the results obtained above as the global dynamics of model (1a)-(1b) in the positive invariant set Ω, we show that the existence of periodic orbits is not possible by Dulac’s criterion [28]:

#### Proposition 9

*Model (1a)-(1b) has no limit cycles or homoclinic loops*

Finally, summarizing all the analysis and propositions of this section, we obtain the global dynamics of model (1a)-(1b):

#### Theorem 1

*Consider the model (1a)-(1b)* 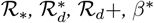, *β*^∗^ *defined above. The following statements hold:*

a. *If* ℛ_∗_ < 1, *the* DFE *is the unique stable equilibrium for* ℛ_*d*_ < 1, *and there exists a unique endemic equilibrium which is stable for* ℛ_*d*_ > 1
b. *If* ℛ_∗_ > 1 *is satisfied and*

1. *if β*_*p*_ *< γ* + *µ, there exist exactly two endemic equilibria* 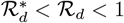, *one of them is stable and competes with the locally stable* DFE, *and one endemic equilibrium for* ℛ_*d*_ > 1.
2. *if* 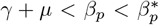, *there exist exactly two endemic equilibria for*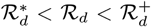, *one of them is stable and competes with the locally stable* EFE, *and one endemic equilibrium for*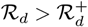.
3. *If* 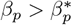, *the* EFE *is the unique stable equilibrium for* 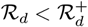, *and there exists a unique endemic equilibrium which is stable for* 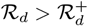.

Figure 4 illustrates the bifurcation diagrams that the model (1a)–(1b) can exhibit according to Theorem 1. When *β*_*p*_ *< γ* + *µ* and condition ℛ^∗^ < 1, Figure 4(a) displays a typical transcritical bifurcation, where the DFE is stable for ℛ_*d*_ < 1. In this case, a unique stable endemic equilibrium emerges for ℛ_*d*_ > 1.

**Figure 4.**
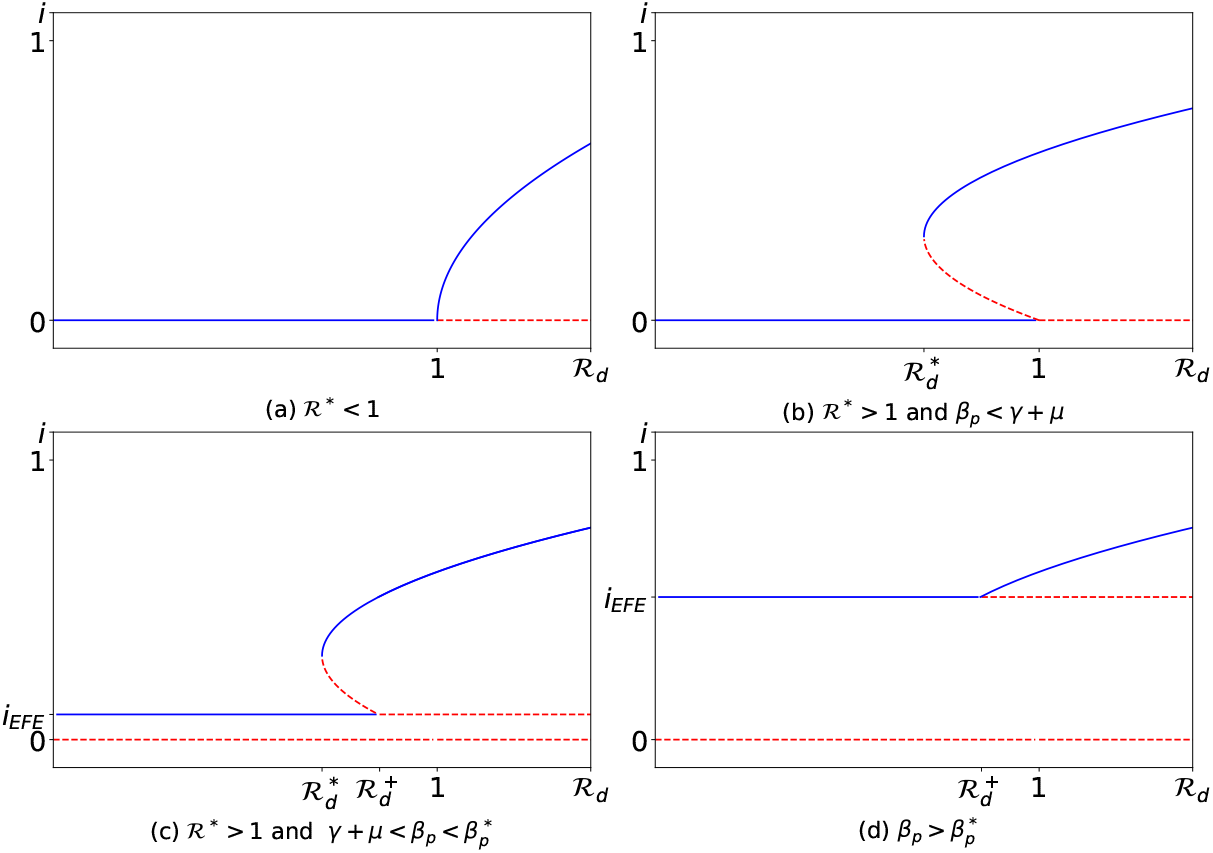
Bifurcation diagrams of model (1a)-(1b) according Theorem 1; blue represent stable equilibrium and red unstable

If *β*_*p*_ *< γ* + *µ* and condition ℛ^∗^ > 1, Figure 4(b) shows the occurrence of a backward bifurcation, characterized by bistability between the DFE and a stable endemic equilibrium for 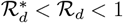.

On the other hand, ℛ^∗^ > 1 and the EFE is feasible in Ω, which is stable for 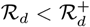. In this setting, Figure 4(c) depicts, for 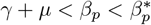, a backward-type bifurcation in which a multistability phenomenon arises between the EFE and the stable endemic equilibrium for 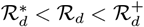.

Finally, Figure 4(d) presents an interesting scenario that resembles a forward bifurcation: the EFE is stable for 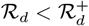, whereas the endemic equilibrium becomes stable for 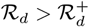 Consequently, no multistability occurs. Nevertheless, we note that the endemic equilibrium emerges for ℛ_*d*_ < 1, which is a distinctive feature of a classical forward bifurcation.

## 4 Discussion

In this work, we study how different modes of transmission interact with a population and influence overall disease dynamics. We develop a model that incorporates both direct (human-to-human) and indirect (human-environment-human) transmission pathways. The coupling between environmental and host dynamics arises from the shedding of pathogens by infected individuals and the subsequent infection of susceptible hosts through contact with contaminated environment. This framework allows us to capture how environmental persistence and human behavior jointly shape the epidemiological outcomes. This framework addresses a critical gap in traditional epidemiological models, which often consider only a single mode of transmission or treat environmental effects as secondary. By explicitly incorporating environmental dynamics into the transmission process, our approach captures complex feedback loops between pathogen persistence, environmental contamination, and host infection. This allows for a more realistic understanding of how outbreaks evolve in settings where sanitation, climate conditions, or pathogen durability play significant roles. Moreover, the coupled formulation can inform control strategies and highlight points of intervention beyond direct person-to-person transmission.

This work extends a previously proposed environmentally mediated transmission framework by incorporating direct human-to-human transmission as an additional infection pathway. While the earlier formulation focused on heterogeneous disease severity and environmental exposure, it did not account for direct transmission between individuals. The present model therefore allows both transmission routes to act simultaneously within a unified theoretical structure [14], leading to significant qualitative changes in the system dynamics. In particular, the analysis reveals multiple dynamical regimes organized by threshold quantities associated with environmental persistence, with the parameter *R*_*d*_ governing the stability of the disease-free equilibrium across broad regions of parameter space.

A central outcome of the analysis is the emergence of multistability for ℛ_∗_ > 1, where ℛ_∗_ represents the expected number of secondary infections produced by an infected individual over its entire infectious period, accounting simultaneously for both direct and environmental transmission routes. In this regime, multiple endemic equilibria may coexist with the disease-free equilibrium, leading to sensitive dependence on initial conditions and abrupt transitions between epidemiological outcomes that are not captured by standard threshold-based interpretations.

An additional theoretical insight concerns the role of the direct transmission parameter *β*_*p*_ as an organizing element of the dynamics. Although *β*_*p*_ naturally defines a classical basic reproduction number ℛ_0_ = *β*_*p*_*/*(*γ* + *µ*), this quantity does not act as a standard epidemic threshold in the present model. Instead, when *β*_*p*_ exceeds *γ* + *µ*, a novel environmental-free equilibrium emerges, enriching the equilibrium structure and giving rise to backward-type bifurcation behavior. This highlights how classical epidemiological quantities may acquire new interpretations in environmentally coupled systems.

From a broader perspective, these results demonstrate how the interplay between direct and environmentally mediated transmission can generate rich and non-intuitive dynamics even in relatively simple models. The presence of multiple stable equilibria and backward-type bifurcation regimes suggests that control strategies targeting a single transmission pathway may be insufficient to guarantee disease elimination. Future work will focus on incorporating more general transmission functions beyond the classical mass-action formulation, such as Holling-type incidence rates, in order to investigate whether additional nonlinear mechanisms further enrich the equilibrium structure and bifurcation scenarios of the model.

## A Proofs of the Propositions

### A.1 Proposition 1

Taking into account the directions of the orbits of the model (1a)-(1b), on the boundary of Ω, it is easy to see that

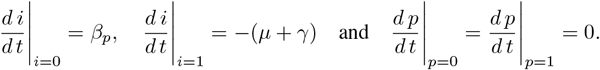

Thus, in every point of the boundary of Ω the vector field of model (1a)-(1b) points to the interior or is an invariant set of Ω. This proves the sentence.

### A.2 Proposition 2

Let us define the function *F* : ℝ^2^ → ℝ^2^, with bifurcation parameter *β*_*p*_, as 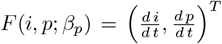 Thus, the jacobian matrix in (0, 0; *γ* + *µ*) is given by:

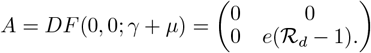

We can see that *A* has a simple eigenvalue *λ* = 0 with right and left eigenvectors given by 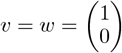, with which we obtain

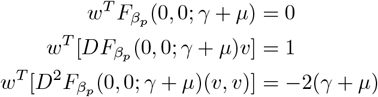

Thus, by Sotomayor’s theorem [20], the Proposition is true.

### A.3 Proposition 3

The Jacobian matrix associated to the *DFE* and the *EFE* are given, respectively, by

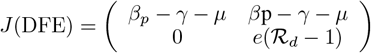

and

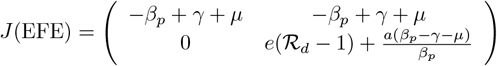

whose eigenvalues are, respectively

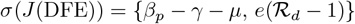

And

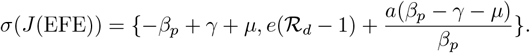

With this, Proposition 3 is evident. In Figure 5, we illustrate both DFE and EFE and their stability for a range of values of ℛ_*d*_; when *β*_*p*_ *< γ* + *µ*, only the first one is biologically feasible and stable for ℛ_*d*_ < 1. However, when *β*_*p*_ *> γ* + *µ*, EFE appears with local asymptotical stability for 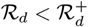 and the DFE becomes unstable for all ℛ_*d*_.

**Figure 5.**
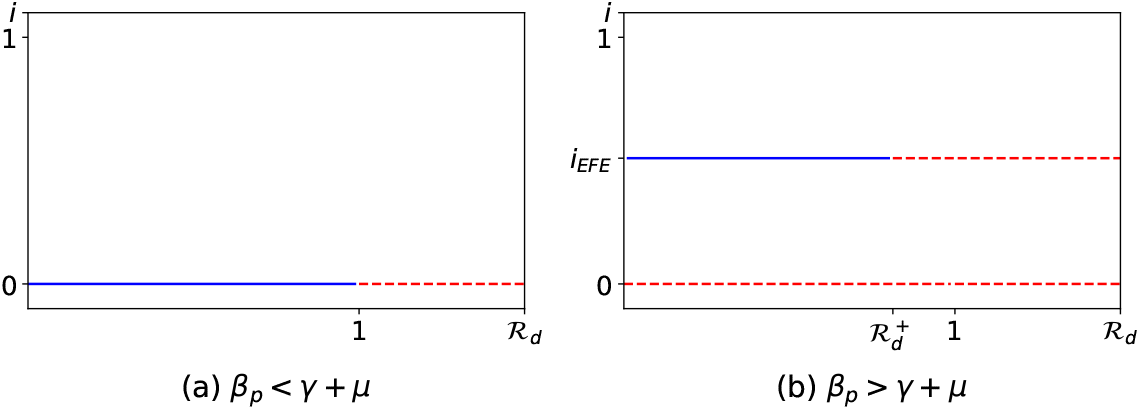
DFE and EFE for some values of *R*_*d*_. Blue represents stable equilibria; red represents unstable equilibria

### A.4 Proposition 4

Evidently, *Q*_3_( ℛ_*d*_) > 0. In addition, observe that if *Q*_2_( ℛ_*d*_) < 0 necessarily implies *Q*_1_(ℛ _*d*_) < 0. With this, Table 1 shows the number of positive roots that the polynomial (10) can have regarding the signs of its coefficients.

**Table 1:** The signal of the possible coefficients of *P*_3_(*i*; _*d*_) and their respective number of positive roots according to the Descartes’ rule.

| $Q_3(\mathcal{R}_d)$ | $Q_2(\mathcal{R}_d)$ | $Q_1(\mathcal{R}_d)$ | $Q_0(\mathcal{R}_d)$ | Number of positive roots |
| --- | --- | --- | --- | --- |
| > 0 | > 0 | > 0 | > 0 | 0 |
|  |  |  | < 0 | 1 |
| > 0 | > 0 | < 0 | > 0 | 2,0 |
|  |  |  | < 0 | 1 |
| > 0 | < 0 | < 0 | > 0 | 2,0 |
|  |  |  | < 0 | 1 |

First, observe that statement (a) is evident by Table 1. To prove the statement (*b*), note that if ℛ_*d*_ = 1, then *Q*_0_(1) = 0 and

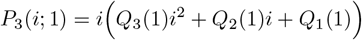

whose roots are 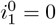, which coincides with the DFE, and the other two roots are given by

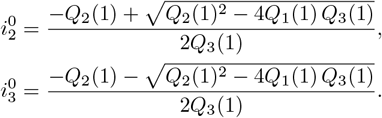

Observe that if we set *Q*_1_(1) < 0, that is, if ℛ^∗^ > 1, then 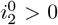 and 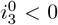. Moreover, let us consider 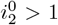, or equivalently

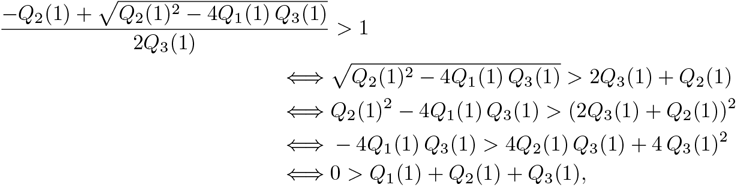

but *Q*_1_(1) + *Q*_2_(1) + *Q*_3_(1) = (*a* + *e*)(*γ* + *µ*) > 0, therefore, 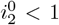, *i*.*e* 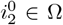. Next, because *P*_3_(*i*; ℛ_0_) is a polynomial, it is a continuously differentiable function and satisfies 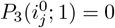, for *j* = {1, 2}, and

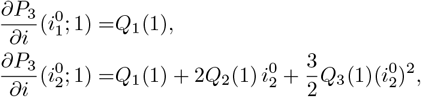

that is, in a generic way, 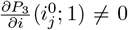, for *j* = {1, 2}. Thus, by the Implicit Function Theorem [19], for each *j* there exists a unique function 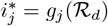 such that *P*_3_(*g*_*j*_(ℛ_*d*_); ℛ_*d*_) = 0 in a neighborhood of 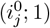, that is, these functions represent the roots of (10) in those neighborhoods. We can obtain the functions *g*_*j*_(ℛ_*d*_) as series expansions around ℛ_*d*_ = 1:

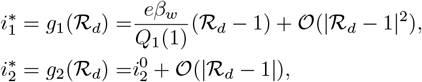

Then, the condition *Q*_1_(1) < 0 guarantees the existence of an open interval when ℛ_*d*_ < 1, such that the roots 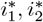 are positive. To determine such interval, note that Δ(1) > 0, then, by continuity, Δ( ℛ_*d*_) > 0 for ℛ_*d*_ < 1. Thus, the interval will be limited by a 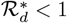 such that 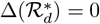. This proves statement (b). Finally, suppose that *Q*_1_(1) > 1, then, we have *Q*_2_(1) > 0 and it is possible to see that 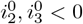, therefore, (10) does not have positive roots, which proves the statement (c).

In Figure 6, we can observe the two scenarios for the roots of the endemic equilibria. Under the condition ℛ^∗^ > 1 there exist two positive roots for 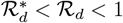 and one root for ℛ_*d*_ > 1, otherwise there exists exactly one positive root for ℛ_*d*_ > 1.

**Figure 6.**
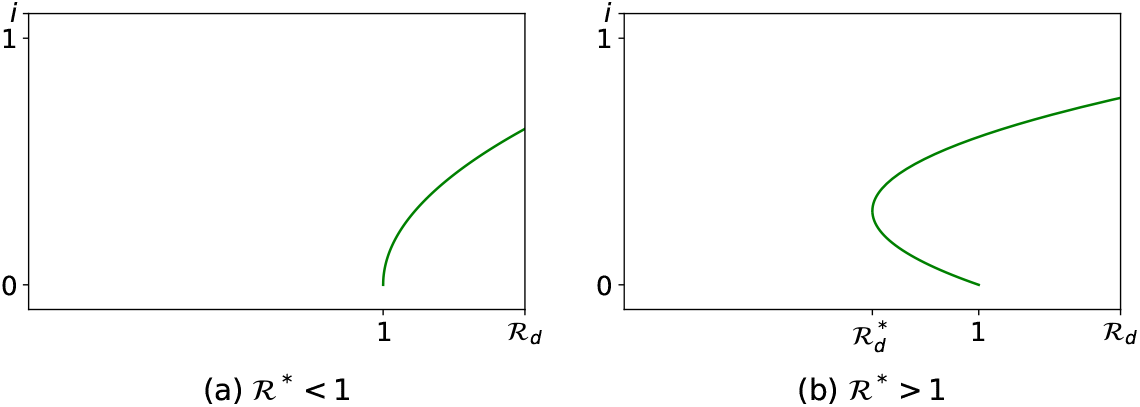
Positive roots of polynomial (10).

### A.5 Proposition 5

Let us define the function *F* : ℝ^2^ → ℝ^2^, with bifurcation parameter ℛ_*d*_,

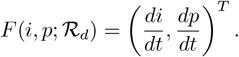

For (*a*), we have

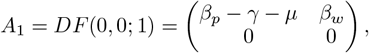

and we can see that *A*_1_ has a simple eigenvalue *λ* = 0 with right and left eigenvectors given by

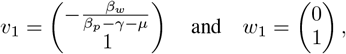

respectively, with which we obtain

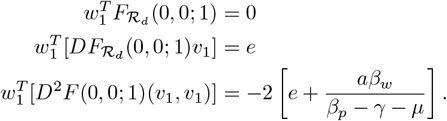

For (*b*), we have

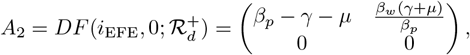

and we can see that *A*_2_ has a simple eigenvalue *λ* = 0 with right and left eigenvectors given by

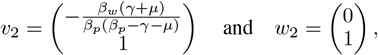

respectively, with which we obtain

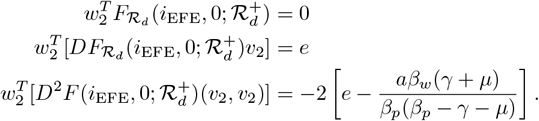

Thus, by Sotomayor’s theorem, the Proposition is true.

### A.6 Proposition 6

The statement (*a*) is obvious from Table 1. For the statement (*b*), consider ℛ^∗^ > 1: if *β*_*p*_ *< γ* + *µ*, then *i*_EFE_ < 0 and, by Proposition 4, the model has two positive roots for 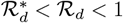 and both are greater than *i*_EFE_ < 0. This proves sentence (*b*) − (1). Next, as we mentioned, if *β*_*p*_ *> γ* + *µ*, then *i*_EFE_ > 0 and, by Proposition 5, occurs a transcritical bifurcation in 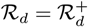, *i*.*e*.,

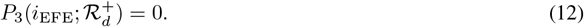

By (8), the roots *i*^∗^ of *P*_3_(*i*; ℛ_*d*_) lowers than *i*_EFE_ are not biologically feasible. This suggests that even when ℛ^∗^ > 1, the phenomenon of multistability is not in Ω as we can see in Figure 4(d). Clearly, this happens when *i*_EFE_ is greater than the vertix of the bifurcation curve. Therefore, it is neccesary to find the value for *β*_*p*_ such that the *i*_EFE_ coincides with the vertex, that is, when *i*_EFE_ is a root of *P*_3_(*i*; ℛ_*d*_) with multiplicity two. For this, as (12) is satisfied, it is possible to find a degree two polynomial for the other two roots of *P*_3_(*i*; ℛ_*d*_), given by

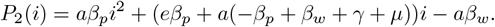

Then, we find the value for *β*_*p*_ such that *P*_2_(*i*_EFE_) = 0, obtaining

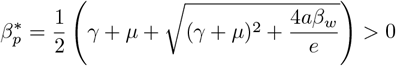

And

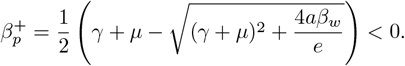

Evidently,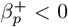then the value 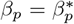causes that *i*_EFE_ coincides with the vertex of the bifurcation curve, and (*b*) − (2) and (*b*) − (3) are true.

### A.7 Proposition 7

Upon implicit differentiation of (10) with respect to ℛ_*d*_, and considering that 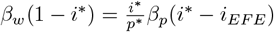 by (7a), we obtain

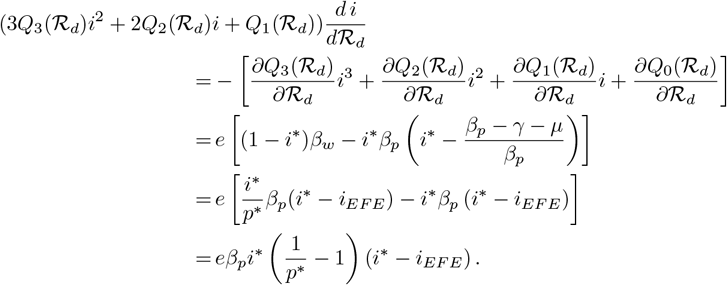

As (*i*^∗^, *p*^∗^) is an endemic equilibrium, by (8), the right hand of the above equation is positive, and the proposition is true.

### A.8 Proposition 8

We compute the jacobian matrix for (*i*^∗^, *p*^∗^) obtaining

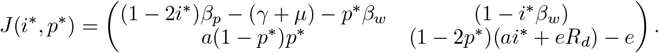

Besides, considering that

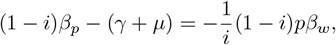

and

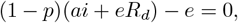

by (7a) and (7b) respectively, we have

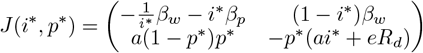

from where, we can obtain

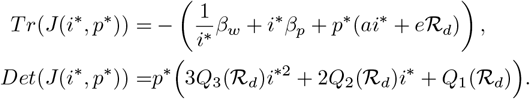

Then, considering the characteristic polynomial of *J* (*i*^∗^, *p*^∗^), given by

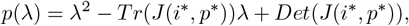

by Routh-Hurwitz’s theorem [1], the Proposition is true.

### A.9 Proposition 9

Consider the function

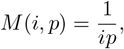

then, we have

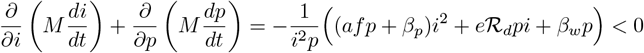

for all *i >* 0, *p >* 0. Thus, by Dulac’s criterion, the sentence is true.

## Notes

### Competing Interest Statement

The authors have declared no competing interest.

